# Cerebellar dopamine D2 receptors regulate preference for social novelty

**DOI:** 10.1101/2019.12.20.884288

**Authors:** Laura Cutando, Emma Puighermanal, Laia Castell, Pauline Tarot, Morgane Belle, Federica Bertaso, Margarita Arango-Lievano, Fabrice Ango, Marcelo Rubinstein, Alain Chédotal, Manuel Mameli, Emmanuel Valjent

## Abstract

The cerebellum, a primary center involved in the control of sensorimotor tasks, also contributes to higher cognitive functions including reward, emotion and social interaction. The regulation of these behaviors has been largely ascribed to the monoaminergic system in limbic regions. However, the contribution of cerebellar dopamine signaling in the modulation of these functions remains largely unknown due to the lack of precise characterization of cerebellar dopaminoceptive neurons. By combining cell type-specific transcriptomic and histological analyses, 3D imaging and electrophysiology we demonstrate that cerebellar dopamine D2 receptors (D2R) in mice are preferentially expressed in Purkinje cells (PCs). While activation of D2R regulate synaptic efficacy onto PCs, their deletion or overexpression in PCs bidirectionally controls preference for social novelty without affecting motor functions. Altogether, these findings demonstrate novel D2R’s roles in PC function and causally link cerebellar D2R levels of expression to social behaviors.

## Introduction

The cerebellum is a brain area that has long been considered to be solely involved in coordination of voluntary movements including the control of posture, balance and gaze orientation^1,2^. However, increasing evidence indicates that the cerebellum also takes part in cognitive and emotional abilities as well as social interactions^3–6^. The control of these various components of motor, cognitive and affective behaviors is largely ascribed to its modular compartmentalization into functional independent units tightly modulated by monoaminergic inputs which signal arousal, salience and valence ^7–9^. Accordingly, the dense plexus of tyrosine hydroxylase (TH)-positive fibers detected in all cerebellar layers and lobules^10^ provides an anatomical substrate to the influence of monoamines to cerebellar neuronal and synaptic activities. Early evidence suggested the presence of dopaminergic innervation in the cerebellum^11–15^. However, its role within this structure remains largely enigmatic, in part because of the lack of and sometimes conflicting information regarding the distribution and the expression level of cerebellar DA receptors, in particular the dopamine D2 receptors (D2R). Thus, although seminal autoradiography and *in situ* hybridization studies revealed the existence of cerebellar D2R binding sites and D2R mRNA^16–19^, the use of the cerebellum as brain reference area in imaging studies ^20,21^ questions the presence of cerebellar D2R. Clarifying this issue is of paramount importance to determine whether altered D2R expression leads to cerebellar dysfunctions frequently associated with motor, cognitive and social deficits observed in autism spectrum disorders (ASD), bipolar mood disorder, unipolar depression, posttraumatic stress disorder or schizophrenia^22-31^.

In this study, we combined cell type-specific molecular and imaging analyses, electrophysiology and mouse behavior to parse the role of D2R in the cerebellum. We demonstrate that D2R are widely distributed throughout the cerebellar cortex (CC) and preferentially expressed by Purkinje cells (PCs). We show that D2R are functional in adult mice and regulate excitatory synaptic efficacy onto PCs. Using adenoviral approaches to selectively delete or overexpress D2R in PCs, we reveal that cerebellar D2R play a key role modulating the preference for social novelty without affecting the motor performance and the coordination of voluntary movements. Together, our findings unveil an unsuspected role of D2R in the cerebellum in the regulation of social interaction and may have important implications in the understanding of the etiology of socially-related disorders.

## Results

### Distribution of D2R in cerebellar neurons

The lack of sensitive tools to detect low expression levels of D2R in the brain largely hampered the precise characterization of cerebellar D2R-containing cells. To determine their identity, we used BAC transgenic mice expressing fluorescent reporter protein under the control of *Drd2* promotor^32^. We first examined the expression pattern of GFP-positive cells in the cerebellum of *D2-RCE* mice. The analysis of GFP immunofluorescence allowed the identification of D2R-positive cells in the CC (Figure 1a_1-_a_2_). A dense GFP staining was observed in the deep cerebellar nuclei (DCN) (Figures 1a_1_ and S1a) most likely corresponding to the terminals of PCs identified at higher magnification (Figure 1a_2_, a_3_). To gain insights into the cellular characterization, we next examined the distribution of the cerebellar D2R-expressing cells in the *D2-RiboTag* mice which express the ribosomal protein Rpl22 tagged with hemagglutinin (HA) selectively in D2R-containing cells^32^. The analysis of HA-immunoreactivity in cerebellar slices (Figures 1b and S1b) or in cleared whole cerebellum using the iDISCO method^33^ (Video S1) confirmed that cerebellar D2R neurons were preferentially localized in the CC (Figure 1b and S1b). Although discontinuous pattern of GFP- or HA-positive labelled PCs were observed, they did not match the expression pattern of zebrin II, a well-established marker of PCs compartmentalization (Figure 1c). We also identified few D2R-expressing neurons in the molecular and granular layers (Figure 1b_2-_b_3_) as well as in the fastigial and interpositus nuclei (Figure S1b) suggesting that D2R were not restricted to PC layer. To further broaden our analysis, we determined the distribution and the percent of HA-positive cells in the lateral cerebellum (Simple lobule, Crus I, Crus II and Paramedian lobule) and in the cerebellar vermis (from lobule II to X) throughout the three cerebellar layers (Figures 1d-1e, S2, Table S1 and S2). Our quantification revealed that a high proportion (between 87% and 96%) of CC HA-positive cells were localized in PC layer compared to the molecular layer (between 3% and 12%) while only few HA-expressing cells were detected in the granular layer (Figure 1e, S2 and Tables S1 and S2). Altogether, these results indicate that D2R are mostly expressed in the PC layer, with homogenous distribution throughout the cerebellar vermis and hemispheres as well as among lobules.

**Figure 1.**
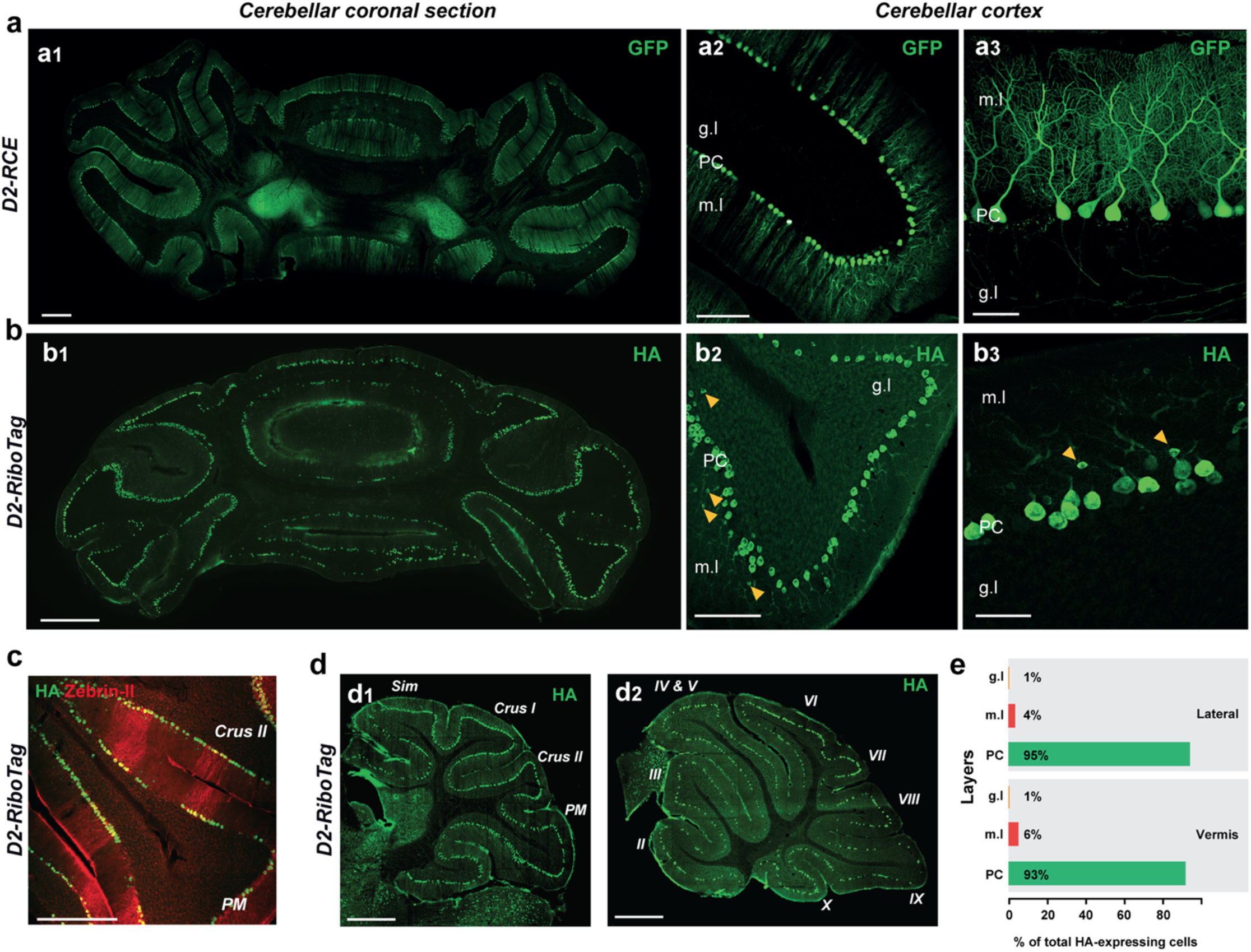

### Dopamine D2R are preferentially expressed by Purkinje cells

After manual dissection of the CC (Figure S3), we next characterized the molecular identity of cerebellar D2R-positive cells in the CC by using the RiboTag methodology^34^. The enrichment of *Drd2* and several gene products known to be expressed in PCs (*Pcp2, Gad1, Slc32a1, Pvalb* and *Calb1*) after HA-immunoprecipitation of CC extracts (pellet fraction) indicated the presence of mRNA encoding the D2R in PCs (Figure 2a-b). Double immunofluorescence analysis performed in *D2-RiboTag* and *D2-RCE* mice confirmed that a high proportion of HA- and GFP-labeled cells co-localized with PCs markers including DARPP-32, G-substrate, calbindin-28kD (CB) and parvalbumin (PV) (Figures 2c-d, and S4). Intriguingly, HA-positive cells do not co-localized with calretinin (CR), a protein known to be expressed in cerebellar Granule cells (GrCs)^35^. The de-enrichment in *Calb2* observed after HA-immunoprecipitation further confirm the absence of D2R in GrCs (Figures 2b, S5a). The enrichment of *Accn1* and *Nos1ap* gene products, two markers of GABAergic molecular layer interneurons (MLI-INs)^36^ as well as the presence of HA/PV cells in the molecular layer (yellow arrowheads) indicated that a fraction of baskets and stellate cells might express D2R (Figure 2b-c). Moreover, the significant enrichment in *Acan* but not in *Grm2* and *Grp* suggested that D2R might also be expressed by a small proportion of Lugaro cells (LCs) but not by Golgi (GoCs) and Unipolar Brush cells (UBCs) (Figure 2b). The presence of *Drd2* mRNA in these distinct neuronal populations was further confirmed by *in situ* hybridization. Indeed, *Drd2* mRNA was found in cells expressing the gene encoding for the vesicular GABA transporter (*Slc32a1*) in the CC including PCs and few MLI-INS (Figure 2e). By contrast, the lack of co-localization of HA and GFP with GFAP (Figure 2f-g) and Iba1 (Figure S5b) as well as the depletion of *Cnp, Itgam* and *Gfap* gene products found after HA-immunoprecipitation, indicated that D2R were not expressed in oligodendrocytes, microglia and glial cells (Figure S5c). This latter observation was confirmed by using *Gfap-RiboTag* mice in which no differences in *Drd2* mRNAs levels were observed between the pellet and the input fractions (Figures S5d-e). We next determined the relative abundance of the two major D2R isoforms generated by alternative splicing, namely the D2R long (D2RL) and D2R short isoform (D2RS)^37,38^, known to display distinct pre and postsynaptic localization within the neuron^39^. To do so, we performed qRT-PCR analysis after HA-immunoprecipitation using designed primers including the sequence of the exon 6 deleted or not by alternative splicing. Two bands corresponding to both *Drd2* isoforms (*Drd2L* ∼250 bp and *Drd2S* ∼160 bp) were detected (Figure 2h). However, the high intensity of the band identifying the *Drd2L* isoform suggested that D2R were mainly expressed at postsynaptic compartments (Figure 2h). Finally, western blot analysis also indicated the presence of the D2R protein in the CC extracts of wild type mice (Figure 2i). Altogether, our results indicate that D2R are preferentially expressed by PCs and to a lesser extent by MLI-INs and Lugaro cells.

**Figure 2.**
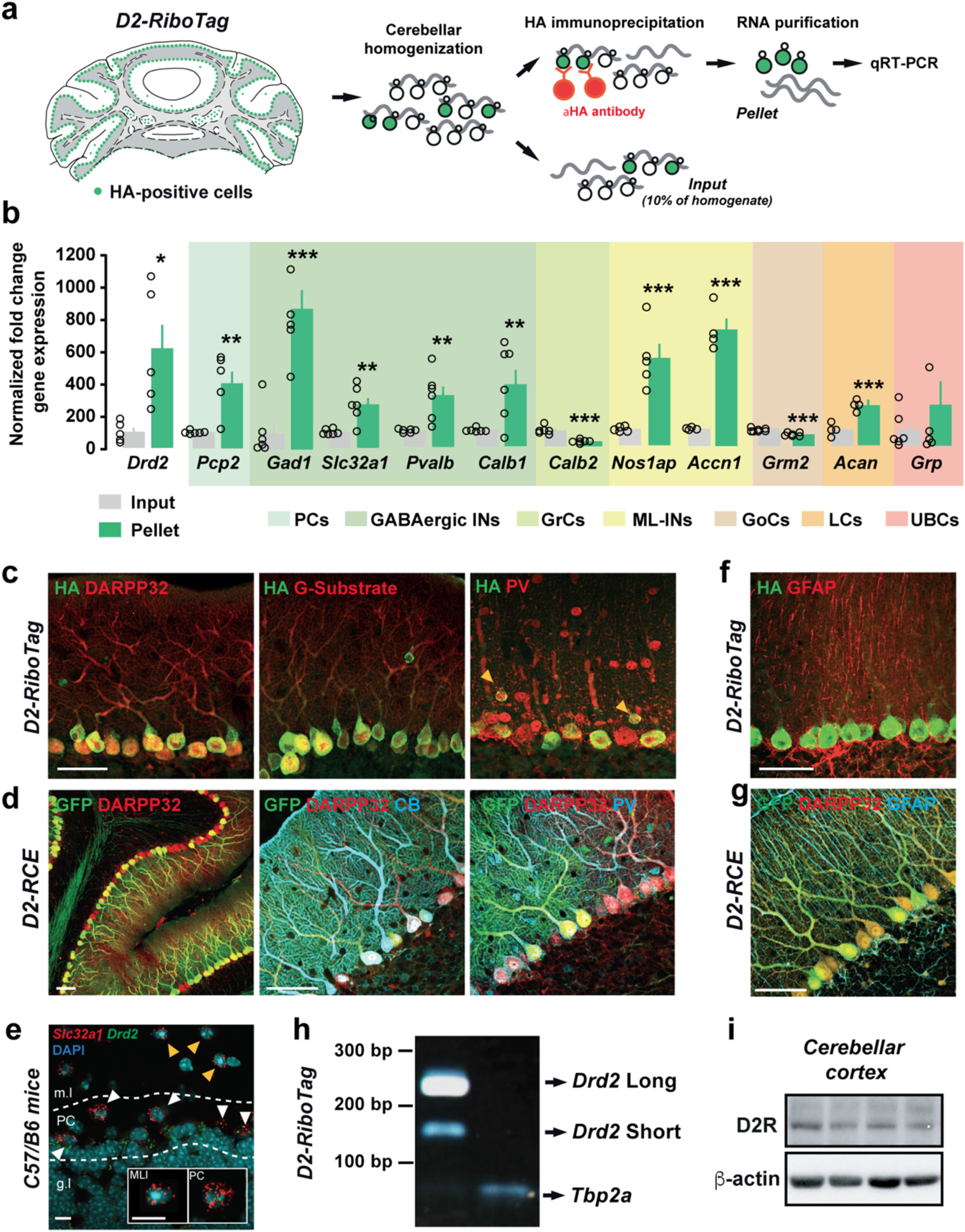

### Activation of D2R modulate excitatory transmission in Purkinje cells

Activation of D2R decreases AMPA-evoked responses in the striatum^40^. However, the effects of D2R activation on PCs AMPA-mediated transmission remain unknown. We first assessed the state of phosphorylation of the AMPA subunit GluA2 at S880 site (pS880-GluA2) –as a biochemical correlate of PCs excitability– following D2R activation (Figure S6a)^41,42^. Western blot analysis performed on cerebellar slices revealed a decrease of pS880-GluA2 (22% ± 8%) after bath application of the D2R agonist quinpirole (10 μm), without modifying the total expression of the receptor (Figures 3a-b and S6b). This transient effect, observed 15 min after quinpirole application, suggested that direct D2R activation might decrease synaptic excitation onto PCs. To directly test this hypothesis, we recorded Parallel Fiber-evoked PC excitatory postsynaptic currents using cerebellar slices of wild type mice. Bath application of quinpirole (10 µm) during 5 min produced a transient reduction of the PF-PC EPSCs amplitude (20.6 ± 4.5%) compared to the initial baseline (Figure 3c). This decrease was independent of neurotransmitter release probability changes since no difference was found in the paired-pulse ratio facilitation measured at PF-PC synapses after application of quinpirole (Figure 3d). Together, these results suggest that D2R expressed at PCs are functional and modulate synaptic excitation onto PCs.

**Figure 3.**
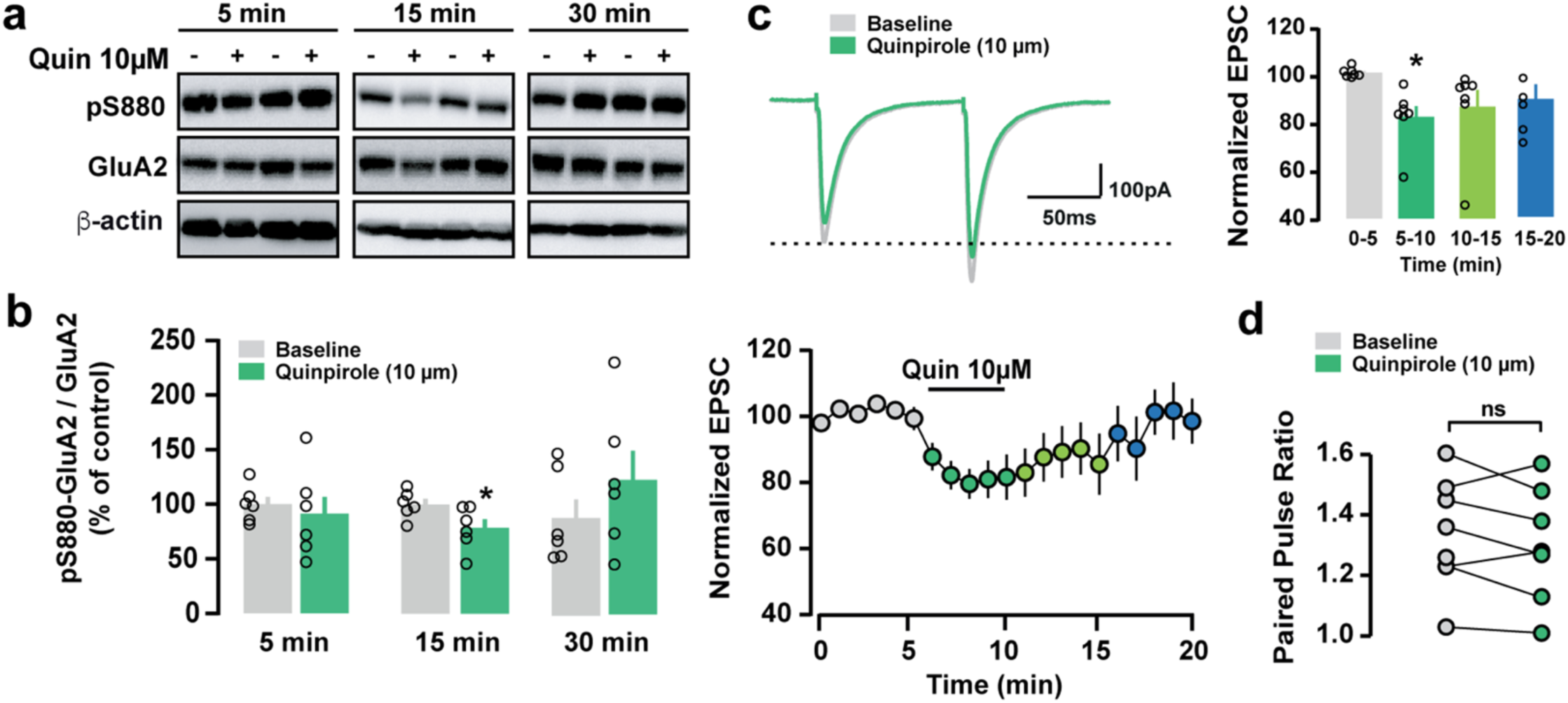

### Generation and validation of conditional D2R knockout mice in Purkinje cells

To investigate the role of D2R expressed in PCs, we generated conditional PCs D2R knockout mice (PC-D2R-cKO). To do so, adeno-associated viral vectors expressing Cre-recombinase selectively in PCs were injected into the CC of *Drd2f/f and Drd2*+/+ mice. We first used an AAV serotype 8 (AAV8-Cmv-Cre-eGFP) known to be one of the most effective vectors to produce selective and high rate of transgene transduction in PCs^43^ (Figure 4). Because D2R are uniformly expressed throughout the CC, 9 stereotaxic injections of AAV8-Cmv-Cre-eGFP were performed on the CC of *Drd2*+/+ and *Drd2f/f* mice to ensure a large D2R invalidation (Figures 4a and S7). Three weeks after surgery, GFP and Cre expression were evaluated in the cerebellum of injected mice. Immunofluorescence analysis revealed that all eGFP positive cells co-localized with CB and Cre recombinase, thereby confirming the selective transduction in PCs (Figures 4b-c). Western blot analysis of Cre recombinase expression levels indicated that PCs were equally transduced in both control and PC-D2R-cKO mice (Figure 4d). Importantly, qRT-PCR analysis performed on isolated eGFP-positive PCs from mice injected with AAV8-Cmv-Cre-eGFP indicated that the expression of *Drd2* mRNA was strongly decreased in PC-D2R-cKO mice compared with the control group (Figure 4e). The invalidation of D2R was further confirmed by double immunofluorescence analysis performed in cerebellar slices of *Drd2f/f* mice injected with AAV8-Cmv-Cre-eGFP. Indeed, we found that D2R immunoreactivity observed at the PC layer (most likely corresponding to PCs) was strongly reduced in all eGFP-positive neurons (Figure 4f). Finally, the deletion of D2R from PCs was functionally efficient since the transient reduction of the PF-PC EPSCs amplitude induced by quinpirole (22 ± 2.2%) observed in eGFP-negative PCs was prevented in eGFP-positive PCs lacking D2R (Figure 4g-i). Of interest, paired-pulse ratio facilitation measured at PF-PC synapses was unaffected by the deletion of D2R in PCs (Figure 4j). Altogether, these results demonstrate that D2R expressed by PCs are involved in the regulation of their excitability upon D2R-agonist bath application and further validate our strategy to generate selective PC-D2R-cKO mice.

**Figure 4.**
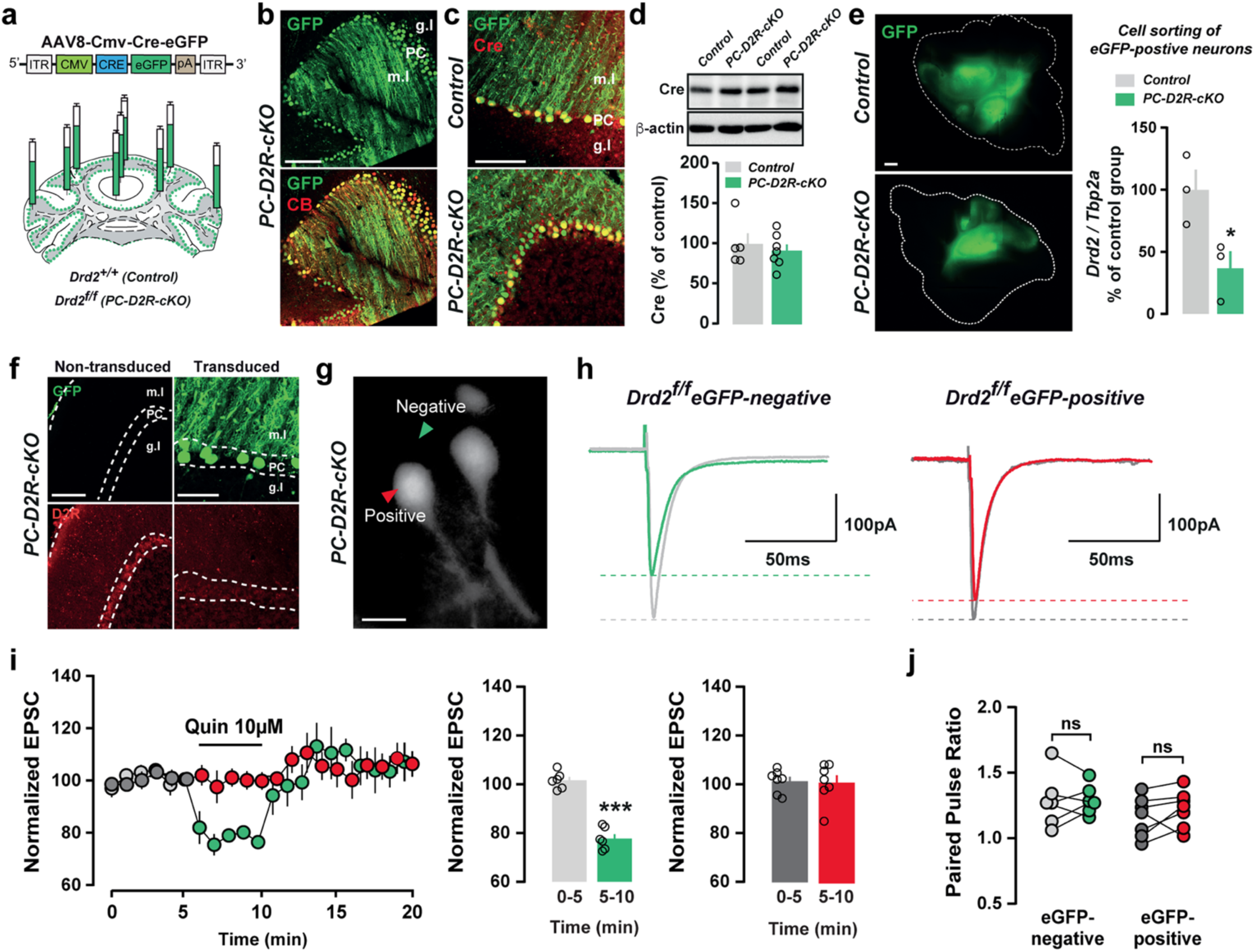

### D2R expressed in Purkinje cells regulate preference for social novelty

Control and coordination of voluntary movements (posture, balance, coordination, gaze orientation) are classically ascribed to PCs functions^1^. Because D2R modulate PCs excitatory transmission, we first assessed the impact of PCs D2R invalidation on locomotion, motor coordination and cerebellar learning. No difference in horizontal and vertical locomotor activity measured in a circular corridor was found between PC-D2R-cKO and control mice (Figure S8a). Moreover, PC-D2R-cKO mice had performance similar to control mice in accelerating rotarod, beam walking and coat-hanger tests, further indicating that motor routines and coordination were not altered in D2R-cKO (Figures S8b-d). In addition to its role in movement control, the cerebellum also takes part in cognitive abilities, regulation of emotions and social interactions^5,6^. We therefore tested whether D2R deletion from PCs might alter social interactions using the 3-chamber test^44^. Sociability was first assessed by measuring the time mice spent exploring an object and an unfamiliar juvenile mouse (stranger 1, S1) (Figures 5a, 5b and S9a). Both control and PC-D2R-cKO mice spent more time interacting with S1 than with the object (Figures 5a, 5b and S9a), a phenotype not due to the preference for one of the chambers towards the other neither for altered locomotor exploration (Figures S10a-e). Although no significant differences were observed between control and PC-D2R-cKO mice on the total time interacting with S1, PC-D2R-cKO mice tend to spend more time sniffing S1 than control mice suggesting that lack of D2R in PCs might enhance social interaction (Figures 5a-b, S9a). We then evaluated preference for novel social stimuli. As expected, control mice spent more time in close interaction with novel intruder (stranger 2, S2) than with the familiar one (S1) (Figure 5c). The interaction time with S2 was significantly increased in PC-D2R-cKO mice, an effect independent of the entries and total time spent in S1 and S2 chambers (Figures 5c and S10f-h). To further characterize the preference for social novelty, the amount of time that the test mouse spent sniffing the novel intruder S2 was evaluated by epochs of 2 minutes. In wild type mice the time in interaction with S2 rapidly declined after 2 min (Figures 5d and S9b). In contrast, PC-D2R-cKO mice spent more time interacting with the new mouse for the entire duration of the 10-min test (Figures 5d and S9b), specially during the first 4 min.

Similar results were obtained using an AAV in which the Cre recombinase is driven selectively in PCs by the minipromoter Ple155(Pcp2) (AAVDJ-Pcp2-Cre)^45^, further confirming that reduced D2R levels in PCs promote preference for social novelty (Figures S11 and S12).

**Figure 5.**
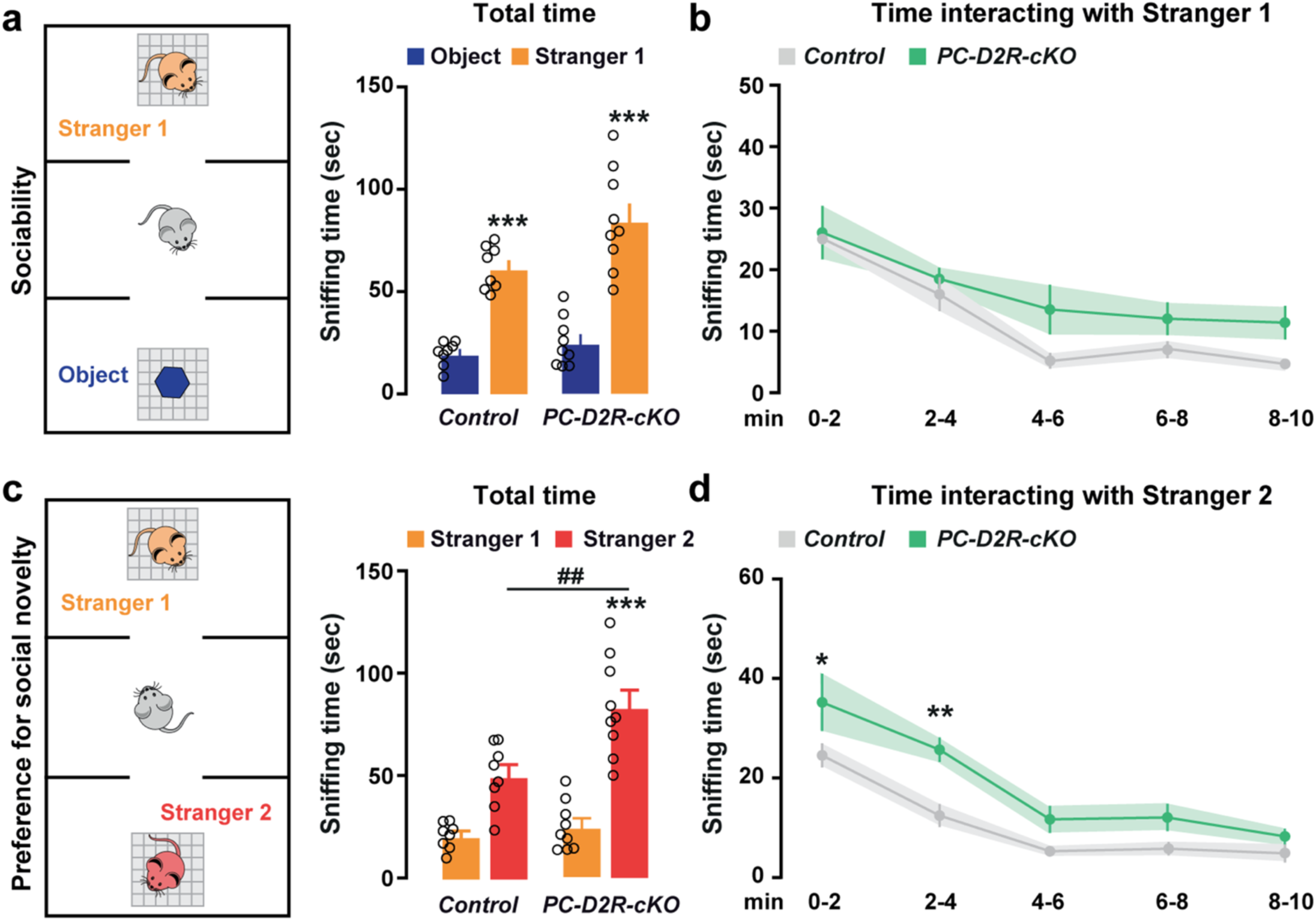

### D2R overexpression in Purkinje cells impairs preference for social novelty

Our previous results suggest that preference for social novelty might be intimately linked to PCs D2R levels. To directly test this hypothesis, enhancement of D2R signaling was achieved by injecting AAV8 virus expressing D2R and eGFP (AAV8-Cmv-mDrd2-IRES-eGFP) in the CC of C57/Bl6 mice (Figure 6a). Double immunolabeling of GFP and PCs markers (PV and CB) confirmed that virally expressed D2R were restricted to PCs (Figure 6b). Moreover, qRT-PCR and western blot analyses showed that AAV8-Cmv-mDrd2-IRES-eGFP injected mice displayed a significant overexpression of *Drd2* mRNAs compared to control mice injected with AAV8-Cmv-eGFP (Figure 6c). Importantly, the differential expression was also confirmed at the protein level by western blot (Figure 6d). Both D2R-overexpressed (PC-D2R-OE) and control mice showed comparable locomotor activity and motor coordination indicating that upregulation of D2R selectively in PCs does not affect cerebellar motor functions (Figure S13a-d). Moreover, sociability was not affected by the overexpression of D2R in PCs since both groups spent equivalent amount of time interacting with S1 during the 10-min session (Figures 6e-f and S14a). In contrast, upregulation of D2R in PCs impaired social preference. Indeed, PC-D2R-OE mice spent significantly less time interacting with the new intruder (S2) compared to control mice (Figures 6g-h and S14b). Of interest, no difference in the total time in the S1 and S2 chambers was found between the two groups (Figure S15). On the other hand, time course of interaction analysis strongly suggested a rapid loss of social interest in PC-D2R-OE mice, especially during the epoch 4 to 6 min of the entire 10-min test (Figures 6h and S14b). Altogether, these results endorse the causal link between cerebellar D2R expression levels and the modulation of social novelty behavior.

**Figure 6.**
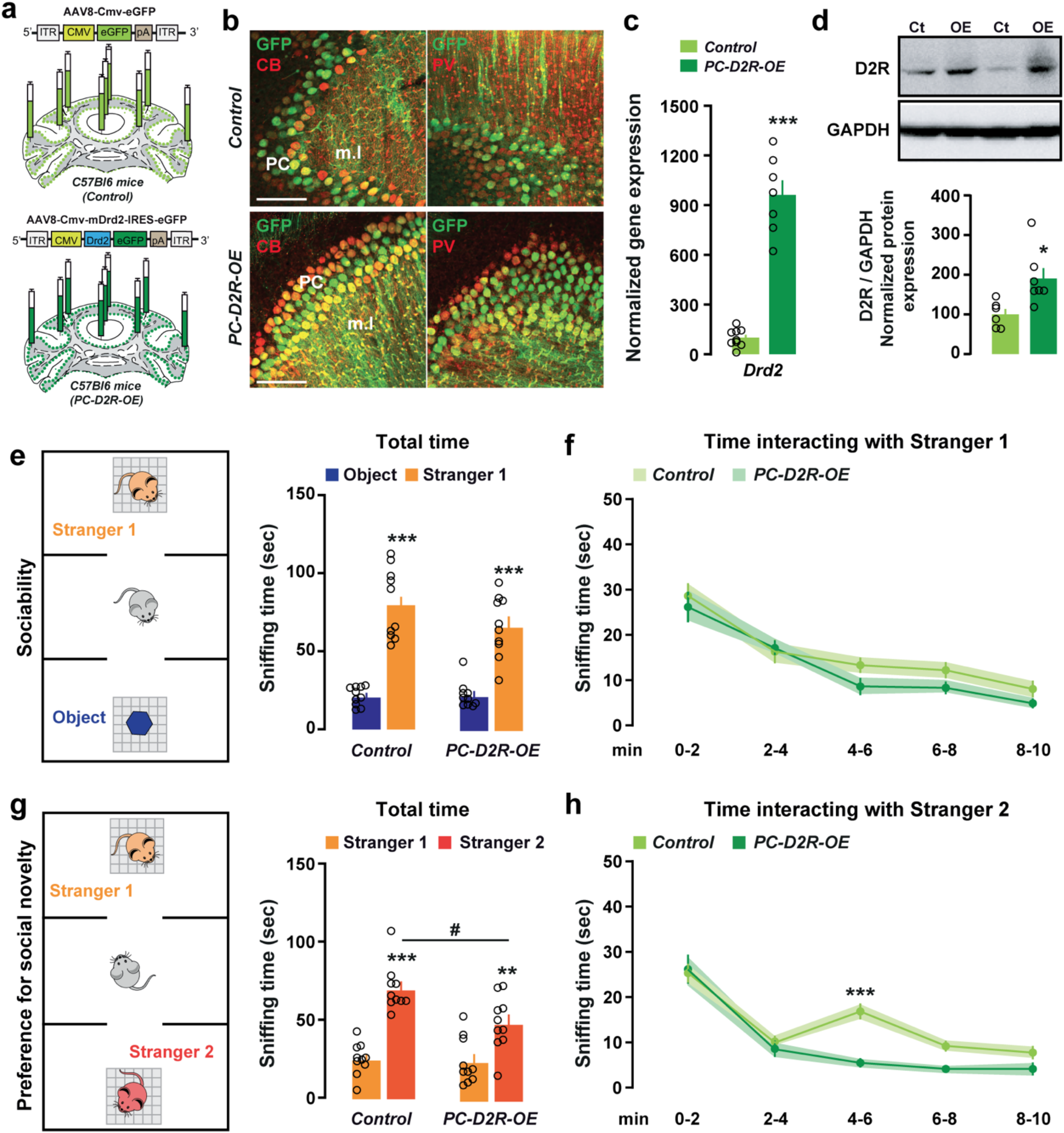

## Discussion

In this study, we demonstrate that cerebellar D2R neurons are homogenously distributed throughout the cerebellar vermis and hemispheres. We establish that D2R are preferentially expressed in PCs in which they contribute to the regulation of the excitatory transmission. We use different adenoviral approaches to selectively delete or overexpress D2R in PCs and demonstrate that D2R expression levels gate the preference for social novelty while having no effect on motor cerebellar functions such as coordination of voluntary movements.

Although several detailed brain mappings of D2R at both mRNA and protein levels were established, contrasting results casted the doubt about the presence of D2R in the cerebellum. In this study, the use of BAC transgenic mice expressing fluorescent protein or tagged-ribosomes was determinant to parse the distribution and identity of cerebellar D2R-containing neurons. Thus, volume imaging of whole-mounted HA immunolabelling unambiguously identified PCs as the main cerebellar cell population expressing D2R. Such findings provide us a rational interpretation of observations made by using autoradiography in which D2R binding sites were detected in the superficial layers of CC^16–19,46–48^. In addition, the Ribotag methodology allowed us to detect the presence of *Drd2* mRNA in PCs and in few ML-INs. This enrichment was confirmed using single molecule fluorescent *in situ* hybridization suggesting that probes or/and hybridization techniques previously used, were most likely not sensitive enough to detect low levels of *Drd2* mRNA expression in a brain area such as the cerebellum^18,49–51^. Despite contrasting results were obtained regarding the cellular localization of D2R^52–54^, we identified functional D2R in the CC. Indeed, D2R activation produced a transient reduction of the PF-PC EPSCs amplitude, paralleled by a rapid decrease of GluA2 phosphorylation at S880 suggesting a role of D2R in the modulation of synaptic excitation onto PCs. Supporting this idea, the slow inward current observed after a brief strong depolarization of PCs is prevented when D2R are blocked^55^. Further studies will determine to which extend D2R might also contribute to the regulation of short- and long-term cerebellar plasticity.

D2R play an essential role in dopamine-regulated motor control^56,57^. Although D2R’s contribution in the regulation of motor behaviors was mainly ascribed to D2R distributed throughout the basal ganglia circuit, our results suggesting the presence of cerebellar D2R questioned their role in the control of posture, balance and coordination of voluntary movements. The generation of temporally-controlled deletion of D2R selectively in PCs (PC-D2R-cKO) revealed that locomotion, fine motor control and cerebellar-learning performance did not appear to be modulated by D2R expressed in PCs, therefore contrasting with the phenotypes described in full or conditional striatal D2R knockout mice^56–58^. However, one cannot exclude that cerebellar D2R might participate to more selective cerebellar-dependent tasks such as the spontaneous eye blink reflex known to be impaired in some neuropsychiatric disorders associated with DA dysfunctions^59,60^. Instead, our data unveiled a role of cerebellar D2R in social behaviors. Thus, PC-D2R-cKO mice spent more time in close interaction with a new unfamiliar mouse than with a familiar one suggesting that reduced D2R levels in PCs may favor novel social contact. This trait did not result from a global social behavioral deficit since the quality of being social measured by comparing the time spent in interaction with an object and a new unfamiliar mouse was similar between control and PC-D2R-cKO mice. On the other hand, mice overexpressing D2R in PCs displayed a selective impairment in the preference for social novelty revealing a bidirectional control of cerebellar D2R over prosocial behaviors. However, it remains to be addressed whether D2R expressed in PCs may also modulate exploratory responses when other types of novel stimuli are presented, or conversely, if they are particularly involved in the modulation of social novelty behavior. Interestingly, growing evidence supports the hypothesis that distinct DA signaling in specific brain areas may interact dynamically to regulate different aspects of social behaviors. Thus, pharmacological blockade of D2R in the prefrontal cortex (PFC) selectively impaired social novelty discrimination^61^, while contrarily, blockade of dopamine D3 receptors (D3R) improved social memory and enhanced social novelty discrimination^61,62^. Finally, a similar enhancement in social recognition was observed following dopamine D1 receptor (D1R) activation in the PFC and the Nucleus Accumbens (NAc)^63^ where pro-social behavior required intact D1R signaling^64^.

Social interaction is a complex behavior that critically relies on the integrity of midbrain DA ventral tegmental area (VTA) neurons ^64–66^. The dissection of neural circuits regulating social interaction indicate that several neural pathways converging onto different VTA DA neurons may regulate distinct components of social behaviors. Thus, excitatory inputs from the DCN to VTA neurons have been identified as an important pathway involved in the control of social reward^6^. More recently, the superior colliculus to VTA-Dorsal Striatum circuit has been proposed to regulate the orientation towards an unfamiliar conspecific mouse while the anterior cortex to VTA-NAc pathway would be involved in the modulation of conspecific interaction without affecting orienting behaviors^67^. Interestingly, our results indicate that the regulation of pro-social behavior relied in part of DA signaling within the CC. Such findings naturally question the source of cerebellar DA that could contribute to these effects. Early tract-tracing studies indicated that midbrain VTA DA neurons project to the cerebellum^12,13^ (but see^68^). VTA DA neurons projecting back to the cerebellum would form a closed-loop allowing a tight and precise modulation of social behaviors. Supporting this hypothesis, VTA DA neurons projecting to the CC innervate predominantly the Crus I lobule^12^, which has been implicated in the control of social behavior and found impaired in patients with ASD^23,69^ Additional possible sources of DA in the CC include a small fraction of PCs expressing TH^70,71^ as well as noradrenergic fibers ascending from the locus coeruleus, which have been previously shown to release DA in the dorsal hippocampus^72^ and the paraventricular thalamic nucleus^73^. Whether these distinct sources of DA might engage cerebellar D2R in the control of movements, cognition and emotions remain to be determined.

In summary our results reveal an unexpected role of cerebellar D2R in the social novelty preference. Future studies are needed to assess the role of cerebellar D2R in the regulation of some specific components of social contact (e.g. the time following behavior or the nose-to-nose contact), as well as the aggressive behavior or the reinforcing properties of social interaction. Whether D2R expressed by PCs may participate to reward processing through the modulation of the DCN-VTA pathway^6^ still remain to be established.

## Materials and Methods

### Animals

This study was conducted using male and female wild-type C57Bl/6j (Charles River), *Drd2-RCE* (abbreviated as *D2-RCE* in the manuscript), *RiboTag::Drd2-Cre* (abbreviated as *D2-RiboTag* in the manuscript), *RiboTag::Gfap-CreERT2* (abbreviated as *Gfap-RiboTag* in the manuscript), *RiboTag::Drd2-floxed* and *Drd2-floxed* mice of 10-15 weeks old. *Drd2*-RCE mice were produced by crossing the hemizygous BAC *Drd2-Cre* mice (C57Bl/6j, founder ER44) generated by GENSAT (Gene Expression Nervous System Atlas) at the Rockefeller University (New York, NY)^74^ with the ***R****26R* ***C****AG-boosted* ***E****GFP:LoxP (RCE:LoxP)* mice as it was previously described^32^. Homozygous female *RiboTag-floxed*^75^ mice from The Jackson Laboratory were crossed with heterozygous male *Drd2-Cre* and tamoxifen-inducible *Gfap-CreERT2* mice to generate *RiboTag::Drd2-Cre* and *RiboTag::Gfap-CreERT2* mice^32^. Similarly, homozygous female *RiboTag:floxed* were crossed with homozygous *Drd2-floxed* mice (from Dr. Marcelo Rubinstein laboratory). First-generation of mice *RiboTag::Drd2-floxed* heterozygous for the *Drd2-floxed* gene (*RiboTag::Drd2-f/*+) were crossed a second time with homozygous *Drd2-floxed* (*Drd2-f/f*) mice to generate *RiboTag::double-Drd2-floxed* mice (*RiboTag::Drd2-f/f)*. For behavioral experiments, wild-type C57Bl/6j, *RiboTag::Drd2-f/f, RiboTag::Drd2*-+/+, *Drd2-f/f* and *Drd2-*+/+ mice were used.

Animals were maintained in a 12-hour light/dark cycle, in stable conditions of temperature and humidity, with food and water ad libitum. All experiments were in accordance with the guidelines of the French Agriculture and Forestry Ministry for handling animals (authorization number/license D34-172-13). All experiments were conducted blindly to the experimenter and mice were arbitrarily assigned to both viral and pharmacological treatments. The number of animals used in each experiment is reported in the figure legends and the in the statistic report. No statistical methods were used to predetermine sample sizes, but our sample sizes are comparable to those generally used in the field.

### Drugs and Treatments

Quinpirole hydrochloride was purchased from Tocris Bioscience (Cat #1061) and tamoxifen (100 mg/kg) from Sigma-Aldrich. Tamoxifen was administered intraperitoneally in a volume of 10 ml/kg and dissolved in sunflower oil/ethanol (10:1) to a final concentration of 10 mg/ml. To induce the Cre expression in the *RiboTag::Gfap-CreERT2* mice, tamoxifen (100 mg / kg) was administered during 3 consecutive days intraperitoneally in a volume of 10 ml/kg.

### Stereotaxic injection into the cerebellum

Surgeries were performed on 10-12 weeks old *RiboTag:: Drd2-f/f, RiboTag:: Drd2-+/+, Drd2-f/f, Drd2-*+/+ mice and wild type mice. Animals were anesthetized with a mixture of ketamine (Imalgene 500, 50 mg/ml, Merial), 0.9% NaCl solution (weight/vol) and xylazine (Rompun 2%, 20 mg/ml, Bayer) (2:2:1, i.p., 0.1 ml/30 g) and mounted on a stereotaxic apparatus. The microinjection needle was connected to a 10 μl Hamilton syringe and filled with 6 μl of the adeno-associated viruses AAV8-Cmv-Cre-eGFP (titer 7.2E+13 GC/ml; Vector Biolabs #7062) (Great Valley Parkway, Malvern, PA), AAV8-Cmv-eGFP (titer 1E+13 GC/ml; Vector Biolabs #7061) (Great Valley Parkway, Malvern, PA), AAV8-Cmv-m-Drd2-IRES-eGFP (titer 5.3E+13 GC/ml; Vector Biolabs; Lot. 180507#25) (Great Valley Parkway, Malvern, PA) and AAVDJ-Pcp2-Cre (produced in our laboratory). Viral vectors were injected at 9 sites of the CC in order to infect the maximal cerebellar surface. At a rate of 100 nl / min, a total volume of 600 nl were injected at each site during 6 min. The injector was left in place for an additional 5 min before it was removed to allow the diffusion of viral particles away from the site of injection. The coordinates used were: (AP: −5.65 mm, ML: 0 mm, DV: −1.7 mm), (AP: −6,12 mm, ML: −1.8 mm, DV: −2 mm), (AP: −6.12 mm, ML: +1.8 mm, DV: −2 mm), (AP: −6.36 mm, ML: −2.5 mm, DV: −2.8), (AP: −6.36 mm, ML: +2.5 mm, DV: −2.8), (AP: −6.72 mm, ML: −1.2 mm, DV: −2), (AP: −6.72 mm, ML: +1.2 mm, DV: −2.4), (AP: −7.08 mm, ML: 0 mm, DV: −2.4), (AP: −7.56 mm, ML: 0 mm, DV: −2.7). Wounds of mice were sealed by suture (Coated-Vicryl Violet 75 cm, 3/8 circle CC-1 13 mm; #JV1012; Phymep). Animals were then returned to their home cages for a recovery period of 21 days before behavioral evaluation.

### AAVDJ-Pcp2-Cre production

For AAVDJ-PCp2-Cre production we followed the protocol previously described^76^. The AAV plasmid with Pcp2 (Ple155) minipromoter driving expression of iCre was used (pEMS1986, plasmid #49117. Addgene). We selected the minipromoter Ple155 because it selectively transduces PCs as it was previously described^45^. The promoter fragment was isolated via restriction with EcoRI and HindIII enzymes. The viral production was performed with the DJ-packaging system (Cell Biolabs).

### Tissue Preparation and Immunofluorescence

Tissue preparation and immunofluorescence were performed as previously described^77^. Mice were rapidly anaesthetized with euthasol (360 mg/kg, i.p., TVM lab, France) and transcardially perfused with 4% (weight/vol) paraformaldehyde in 0.1 M sodium phosphate buffer (pH 7.5). Brains were post-fixed overnight in the same solution and stored at 4°C. Forty-μm thick sections were cut with a vibratome (Leica, France, RRID:SCR_008960) and stored at −20°C in a solution containing 30% (vol/vol) ethylene glycol, 30% (vol/vol) glycerol and 0.1 M sodium phosphate buffer, until they were processed for immunofluorescence. Cerebellar sections were identified using a mouse brain atlas (Franklin and Paxinos, 2007) and sections comprised between −5.80 mm and −7.0 mm from bregma were included in the analysis. Sections were processed as follows: free-floating sections were rinsed three times 10 min in Tris-buffered saline (50 mM Tris-HCL, 150 mM NaCl, pH 7.5). After 20 min incubation in 0.1% (vol/vol) Triton X-100 in TBS, sections were rinsed in TBS again during 10 min and blocked for 1 h in a solution of 3% BSA in TBS. Cerebellar sections were then incubated 72 hours at 4°C with the primary antibodies (Table 1) diluted in a TBS solution containing in 1% BSA and 0.15% Triton X-100. Sections were rinsed three times for 10 min in TBS and incubated for 60 min with goat Cy3-coupled anti-rabbit (1:500, Thermo Fisher Scientific Cat# 10520, RRID:AB_2534029), goat Alexa Fluor 488-coupled anti-chicken (1:500, Thermo Fisher Scientific Cat#A-11039, RRID:AB_2534096), goat Alexa Fluor 488-coupled anti-mouse (1:500, Thermo Fisher Cat#A-11001, RRID:AB_2534069), goat Alexa Fluor 488-coupled anti-rabbit (1:500, Life Technologies Cat#A-11034, RRID:AB_2576217), goat Cy5-coupled anti-mouse (1:500, Thermo Fisher Scientific Cat#A-10524, RRID: AB_2534033), donkey Cy3-conjugated anti-chicken (1:500, Jackson Immunoresearch Cat#03-165-155, RRID:AB_2340363), goat Cy3-coupled anti-mouse (1:500, Jackson Immunoresearch Cat#115-165-003, RRID:AB_2338680) antibodies. Sections were rinsed for 10 minutes (twice) in TBS and twice in Tris-buffer (1 M, pH 7.5) before mounting in DPX (Sigma-Aldrich). Confocal microscopy and image analysis were carried out at the Montpellier RIO Imaging Facility.

Images covering the entire cerebellum and double-labeled images from each region of interest were acquired using sequential laser scanning confocal microscopy (Zeiss LSM780). Photomicrographs were obtained with the following band-pass and long-pass filter setting: alexa fluor 488/Cy2 (band pass filter: 505–530), Cy3 (band pass filter: 560–615) and Cy5 (long-pass filter 650). All parameters were held constant for all sections from the same experiment. Three-four slices per mouse were used in all immunofluorescence analyses (n = 3 mice / staining).

**Table 1:**
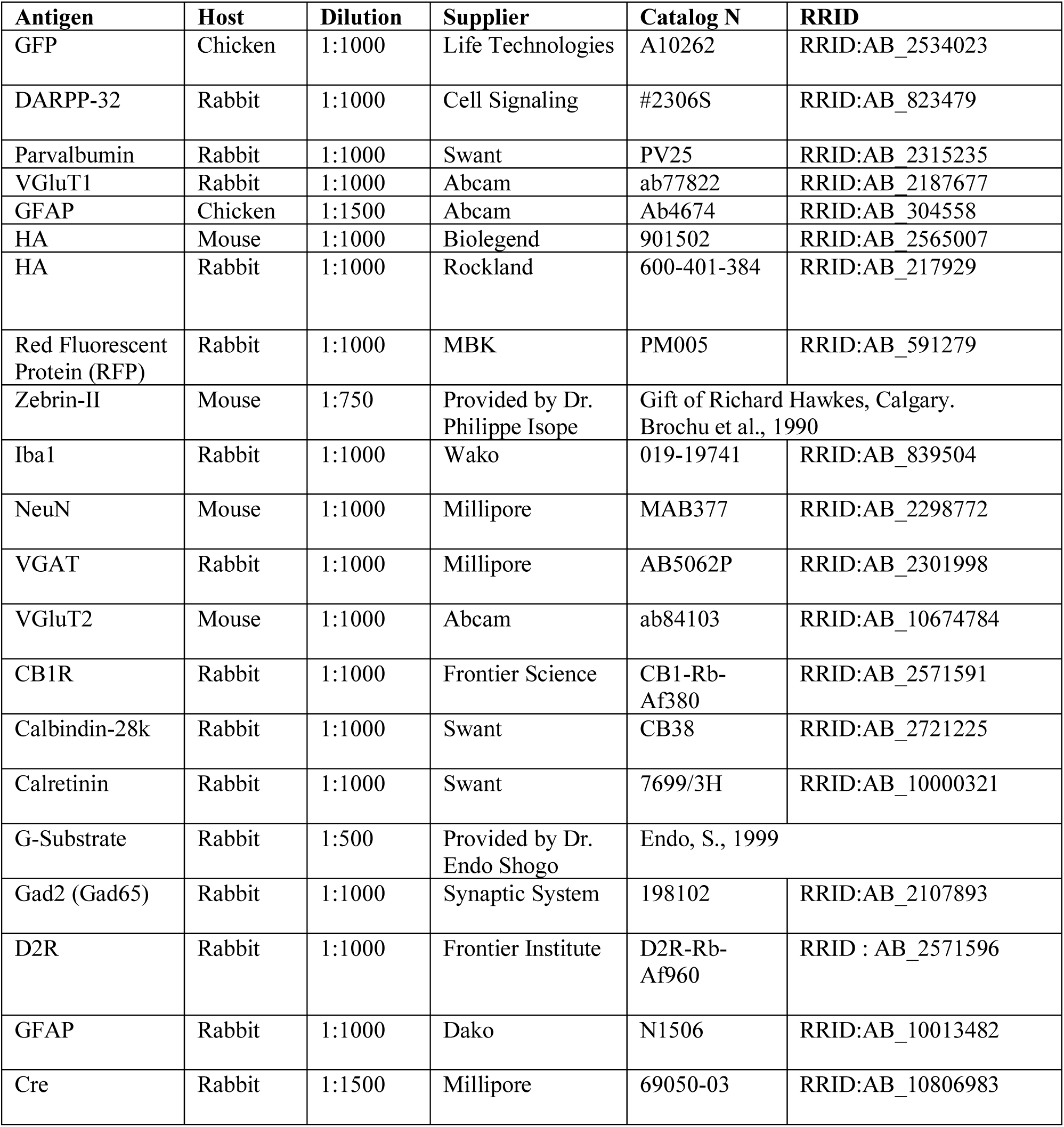
Antibodies used in immunofluorescence.

### Tissue Collection

Mice were killed by cervical dislocation and the heads were immersed in liquid nitrogen for 4 sec. The brains were then removed and sectioned on an aluminum block on ice and the whole cerebellum was rapidly isolated from the stem brain. For the cerebellar cortex and the deep cerebellar nuclei dissection, brains were placed in a matrix and both regions were isolated from ∼1-mm thick coronal section located between 5.80 mm and 6.80 mm posterior to bregma.

### Western Blot

Protein quantification and western blots were performed following the protocol previously described^77^. The whole cerebellum, CC, DCN and cerebellar slices were sonicated in 300 μl, 200 μl and 150 μl and 120 μl of 10% sodium dodecyl sulfate (SDS) and boiled at 100°C for 10 min. In each experiment, samples from different animal groups, treatments or brain regions to be compared were processed in parallel to minimize interassay variations. Following the manufacturer’s instructions, protein contents for each sample were determined by BCA protein assay (Pierce) (Lot# RG235624; Thermo Scientific). Equal amounts of cerebellar lysates were mixed with denaturing 4 Laemmli loading buffer. Samples with equal amounts of total protein (50 μg per lane for Figure 2i and 6d; 25 μg per lane for Figure 3a, S6b, 4d and S11c; 20 μg per lane Figure S3a) were separated in 11% SDS-polyacrylamide gel before electrophoretic transfer onto Immobilon-P membranes (#IPVH00010; Millipore). Membranes were cut horizontally, at different molecular weights to be analyzed with different primary antibodies. Using 4% bovine serum albumin (BSA) in 0.1M PBS, membranes were blocked for 45 min. Next, membranes were incubated for 2h with the primary antibodies (Table 2).

**Table 2:**
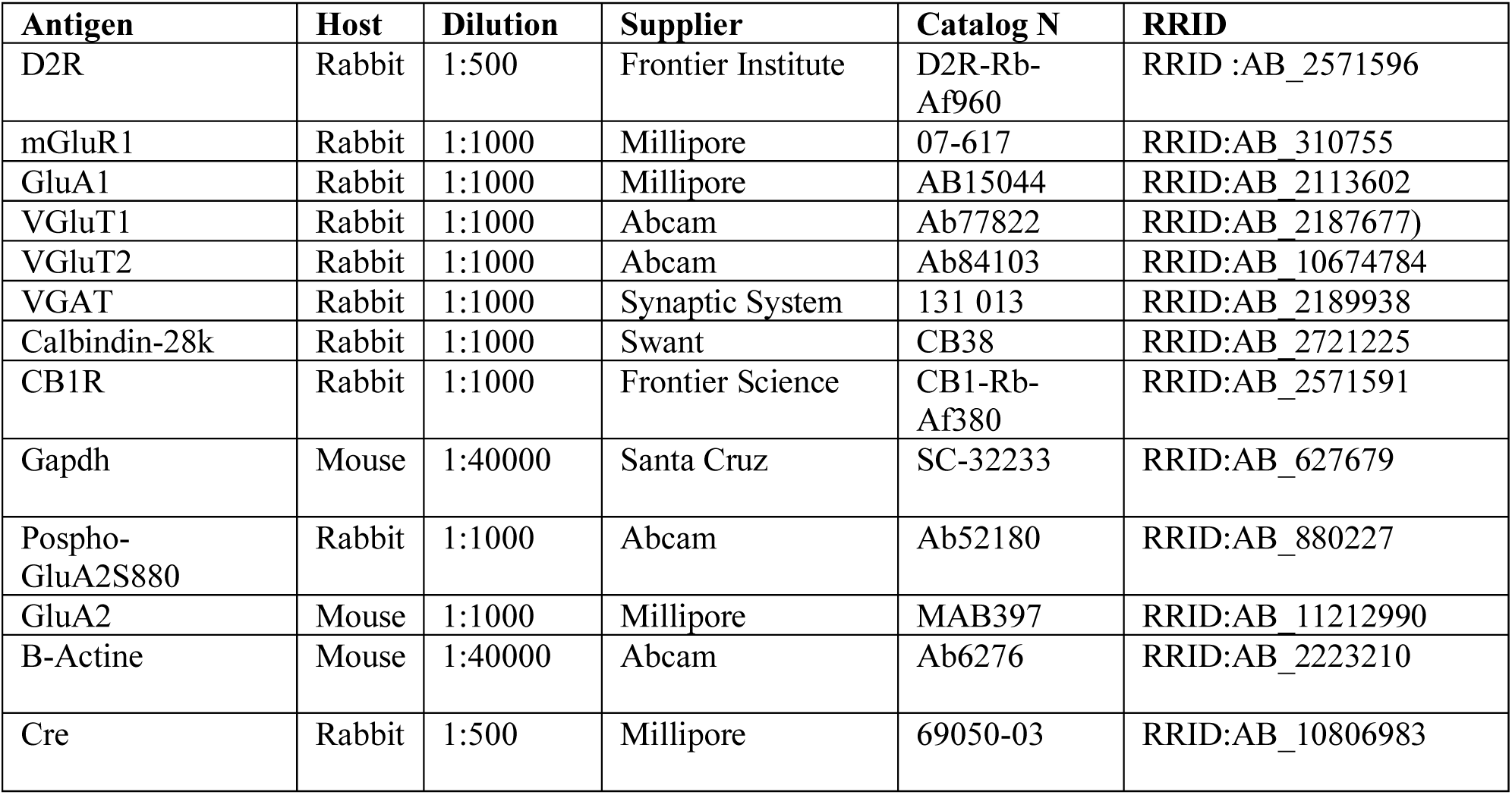
Antibodies used in western blot experiments.

| Antigen | Host | Dilution | Supplier | Catalog N | RRID |
| --- | --- | --- | --- | --- | --- |
| D2R | Rabbit | 1:500 | Frontier Institute | D2R-Rb-Af960 | RRID:AB_2571596 |
| mGluR1 | Rabbit | 1:1000 | Millipore | 07-617 | RRID:AB_310755 |
| GluA1 | Rabbit | 1:1000 | Millipore | AB15044 | RRID:AB_2113602 |
| VGluT1 | Rabbit | 1:1000 | Abcam | Ab77822 | RRID:AB_2187677 |
| VGluT2 | Rabbit | 1:1000 | Abcam | Ab84103 | RRID:AB_10674784 |
| VGAT | Rabbit | 1:1000 | Synaptic System | 131 013 | RRID:AB_2189938 |
| Calbindin-28k | Rabbit | 1:1000 | Swant | CB38 | RRID:AB_2721225 |
| CB1R | Rabbit | 1:1000 | Frontier Science | CB1-Rb-Af380 | RRID:AB_2571591 |
| Gapdh | Mouse | 1:40000 | Santa Cruz | SC-32233 | RRID:AB_627679 |
| Pospho-GluA2S880 | Rabbit | 1:1000 | Abcam | Ab52180 | RRID:AB_880227 |
| GluA2 | Mouse | 1:1000 | Millipore | MAB397 | RRID:AB_11212990 |
| B-Actine | Mouse | 1:40000 | Abcam | Ab6276 | RRID:AB_2223210 |
| Cre | Rabbit | 1:500 | Millipore | 69050-03 | RRID:AB_10806983 |

To detect the primary antibodies, horseradish peroxidase-conjugated antibodies (1:10000) from Cell Signaling Technology to rabbit (Cat# 7074S; RRID: RRID:AB_2099233) or mouse (Cat# 7076S; RRID: AB_330924) were used at and visualized by enhanced chemiluminescence detection (Luminata Forte Western HRP Substrate; Millipore, Cat# WBWF0500). The optical density of the relevant immunoreactive bands was measured after acquisition on a ChemiDoc Touch Imaging System (Bio-Rad) controlled by Image Lab software version 3.0 (Bio-Rad). Representative cropped immunoblots for display were processed with Adobe Illustrator CS6. For quantitative purposes, the optical density values of active phospho-specific antibodies were normalized to the detection of non-phospho-specific antibodies or to β-actin or GAPDH values in the same sample and expressed as a percentage of control treatment or group.

### Cerebellar slices preparation and D2R stimulation by quinpirole

Mice were sacrificed by deep anesthesia with isoflurane inhalation followed by decapitation. The brain was quickly removed and immersed in ice-cold ACSF containing the following (in mm): 126 NaCl, 3 KCl, 1.25 NaH2PO4, 1.3 MgSO4, 26 NaHCO3, 2.5 CaCl2, and 10 glucose equilibrated with 95% O2 plus 5% CO2. Cerebellar sagittal slices (300 µm-thick) were cut by using a vibratome (Campden Instruments, UK) and pooled in 10 ml of ice-cold ACSF buffer. All the solutions used in this experiment were continuously bubbled with 95% O2 and 5% CO2. Once the slices were prepared, they were transferred individually in 6 ml of cold oxygenated ACSF buffer. Cerebellar slices were incubated for 5 min, 15 min or 30 min in ACSF mixed with 6 μl of either quinpirole hydrochloride (Tocris; Cat# 1061, to a final concentration of 10 μM) or distilled water (control). Afterwards, slices were sonicated with 120 μl of 10% sodium dodecyl sulfate (SDS) and boiled at 100°C for 10 min to perform Western blot analysis as described above.

### Polyribosome Immunoprecipitation and RNA extraction

HA-tagged-ribosome immunoprecipitation in the whole cerebellum and CC of *RiboTag::Drd2-Cre* mice was performed as it was previously described^78^ using anti-HA antibody (5ul/sample; Biolegend; Cat#901502) and magnetic beads (Invitrogen, #100.04D). Total RNA contained in the pellet fraction was extracted from ribosome-mRNA complexes using RNeasy Microkit (Qiagen; Cat#74004) and from the input fraction using the RNAeasy Minikit (Qiagen; Cat#74104) following manufacturer’s instructions. Quality and quantity of RNA samples were both assessed using the Nanodrop 1000 spectrophotometer. Between 4 and 6 biological replicates, each one composed of a pool of 2 mice, were used for qRT-PCR analysis.

For the RNA extraction from the CC and DCN (Figure S3c) we used the RNAeasy Minikit (Qiagen; Cat#74104) following manufacturer’s instructions.

### cDNA Synthesis and Quantitative Real-Time PCR

After RNA extraction from pellet and the input fractions of CC *RiboTag::Drd2-Cre* mice, synthesis of cDNA was performed as it was previously described^34^ by using the SuperScript VILO cDNA synthesis kit (Invitrogen) in one cycle program consisting on 10 min at 25°C, 60 min et 42°C, 5 min at 25°C and with a final extension period of 5min at 4°C. Resulting cDNA was used for quantitative real-time PCR (qRT-PCR), using SYBR Green PCR master mix on the LC480 Real-Time PCR System (Roche) and the primer sequences indicated in Table 3. Analysis was performed using LightCycler 480 Software (Roche). The immunoprecipitated RNA samples (pellet) were compared to the input samples in each case. Results are presented as % of change of the pellet fraction versus the input for each gene and normalized to the housekeeping gene *Tbp2a*. For the analysis of the genes expressed in the cerebellar cortex and deep cerebellar nulei, results are presented as linearized Ct-values normalized to housekeeping *Tbp2a* and the ΔΔCt method was used to give the fold change. 4-6 biological replicates are used in these experiments.

**Table 3.**
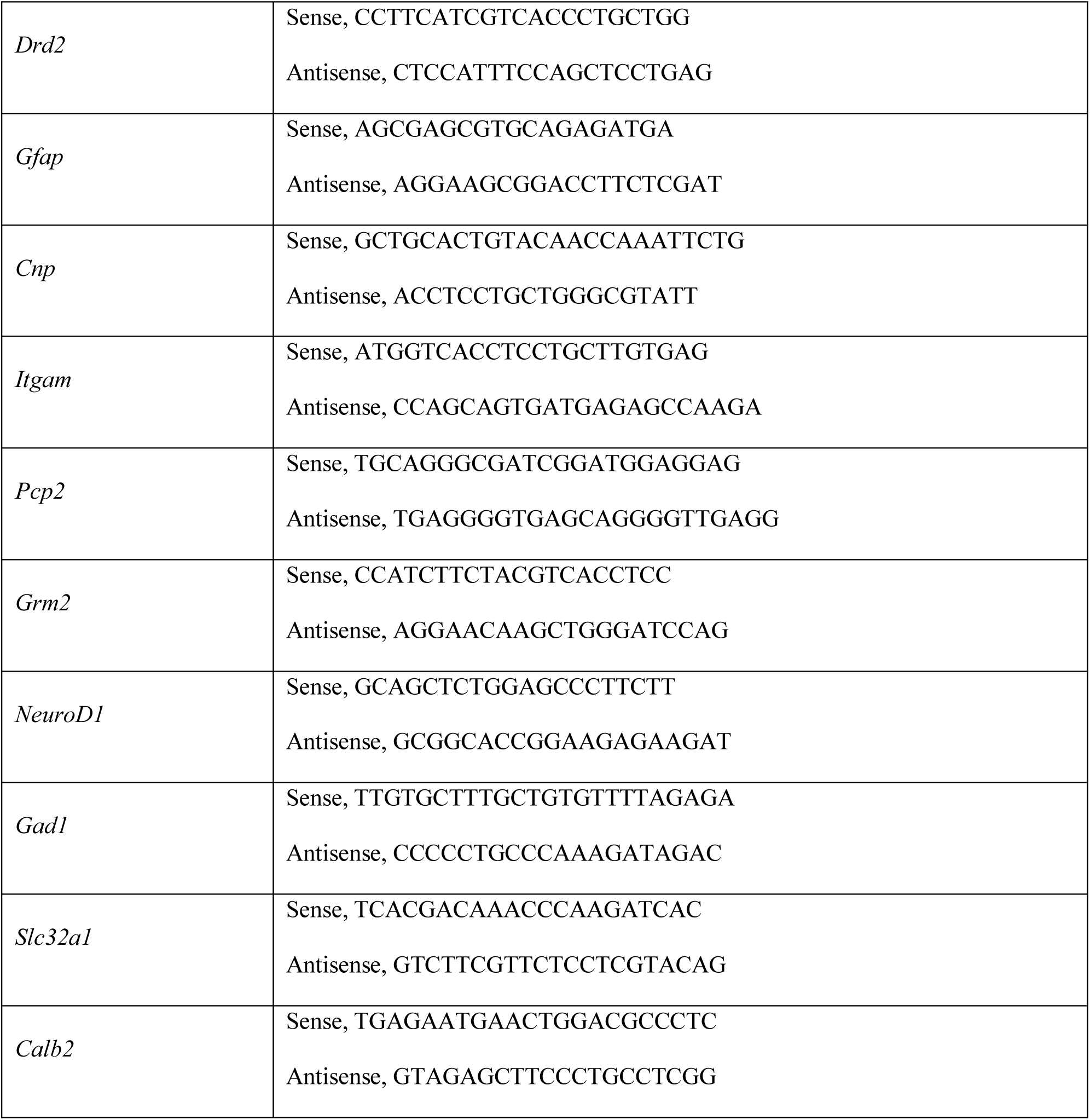
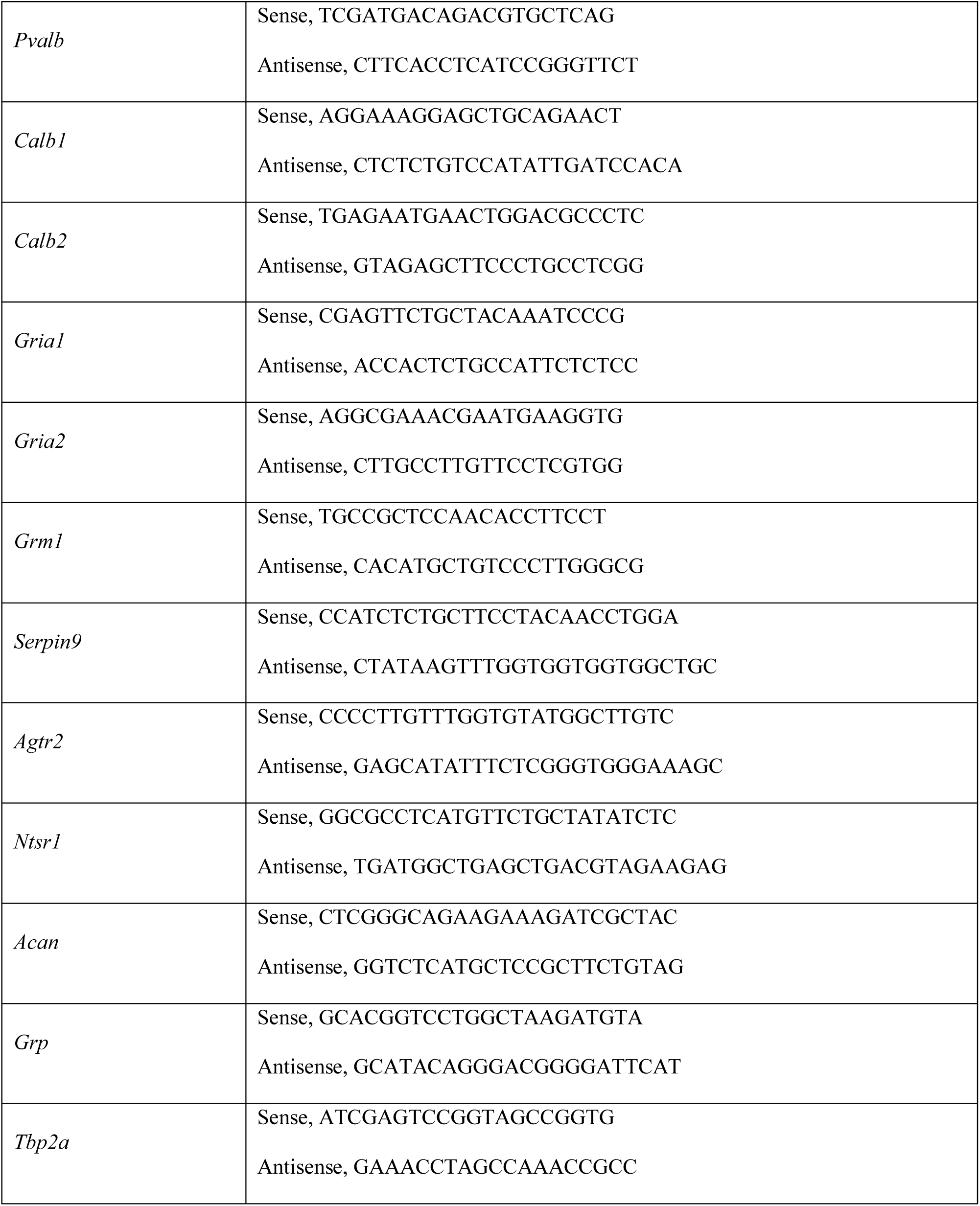
Sequences of primers.

To determine which *Drd2* isoform is the most abundant in the CC, primers were designed to recognize the presence or absence of exon 6 (Table 3). After the qRT-PCR analysis, pellet samples were collected from the qRT-PCR plate and mixed with Purple Gel Loading Dye (BioLabs, Cat#B7025S) to be load on 1% agarose gel. Two band were detected in the gel, the upper band (245bp) corresponds with the long isoform of *Drd2* gene, whereas the lower band (158 bp) corresponds to short isoform of *Drd2* gene.

### Cell Sorting

The brains of *Drd2*+/+ and *Drd2f/f* mice injected with AAV8-Cmv-Cre-eGFP viral vectors were used to purify PCs based on enhanced green fluorescent protein (eGFP) expression as previously described^79^. The brains were dissected in frozen ACSF containing 20 mM glucose and 2 mM Cacl2, then cut in 350-µm sections with a chopper. Slices were separated under the microscope and incubated with oxygenation in ACSF containing 20 mM glucose, 2 mM Cacl2, 0.1 µM TTX, 0.05 mM APV, and 0.02 mM DNQX for 1 h. Slices were transferred in ACSF/glucose/Cacl2/pronase (Sigma, P5147). The slices were incubated in 0.5mg/ml of pronase during 40 min. After an hour of new ACSF/glucose/Cacl2/TTX/APV/DNQX incubation, slices were transferred to an ACSF/glucose/Cacl2/1 % SVF medium. Two lobules infected by the viral vectors were micro-dissected, enzymatically and mechanically dissociated, and plated on a 100-mm Sylgard-bottom Petri dish. After the cells settled, fluorescent neurons were aspirated into a micropipette broken to a diameter of 30 to 50 μm. Then, the cells were transferred into a new Sylgard-bottom dish, and the same procedure was applied three times to obtain near full purity of the fluorescent cells. After the third round of purification, around 40-50 cells were transferred into a glassbottom dish in a small drop (1 to 5 µl) where they could be better inspected for purity under a fluorescence compound microscope. Pure samples were transferred in a 500µl-eppendorff tube and immediately lysed for RNA extraction using the RNeasy Microkit (Qiagen; Cat#74004) following manufacturer’s instructions. cDNA Synthesis and qRT-PCR were performed as previously described.

### Single Molecule Fluorescent In Situ Hybridization RNAscope

For the examination of targeted RNA within intact cells, in situ hybridization RNAscope technology was used following the protocol described by the supplier. Mice were decapitated and brain extracted. Tissue was frozen immediately on dry ice and stored at −80°C. Brains were sectioned at −17°C with the cryostat at 14 µm and mounted onto Superfrost Utra Plus slides (Thermo Scientific; Cat# J4800AMNZ). Sections of brain were collected for the whole cerebellum from bregma −5.80 mm and −6.80 mm. Probes for *Drd2* (ACDBio; Cat# 406501-C3) and *Slc32a1* (ACDBio; Cat#319191-C2) were used with the RNAscope Fluorescent Multiplex Kit (ACDBio; Cat# 320850) as described by the supplier. Slides were counterstained for DAPI and mounted with ProLong Diamond Antifade mountant (Invitrogen; Cat# P36961).

### iDISCO

iDISCO methodology was performed in a cerebellum of *RiboTag::Drd2-Cre* mouse following the protocol previously described^33,80^. Primary antibody anti-HA (Biolegend, Cat#901502, RRID:AB_2565007) and secondary antibody goat Alexa Fluor 488-coupled anti-mouse (1:500, Thermo Fisher Cat#A-11001, RRID:AB_2534069) were used. 3D imaging acquisition was performed by using an ultramicroscope I (LaVision BioTec) with the ImspectorPro software (LaVision BioTec) and movies were generated using Imaris x64 software (version 8.0.1, Bitplane).

### Whole-cell patch-clamp electrophysiological recordings

Unless indicated, all chemicals were from Hello Bio (Bristol, UK). Experiments were performed in parasagittal vermis cerebellar slices, which were prepared in ∼2 months old mice as described previously. Artificial CSF (ACSF) contained the following (in mm): 126 NaCl, 3 KCl, 1.25 NaH2PO4, 1.3 MgSO4, 26 NaHCO3, 2.5 CaCl2, and 10 glucose equilibrated with 95% O2 plus 5% CO2. After a recovery time of ≥80 min, slices were transferred to a chamber perfused with ACSF containing 100 µm picrotoxin at a rate of 2–3 ml/min. When indicated, quinpirole hydrochloride (Trocris Bioscience; Cat #1061) was added to the ACSF at final 10 μM concentration. Whole-cell patch-clamp electrophysiological recordings were performed under infrared-differential interference contrast microscopy at 31–32°C with Multiclamp 700B amplifiers (Molecular Devices, Sunnyvale, CA). The location of the stimulating electrode was gently adjusted around the dendritic tree of the patched cell. Patch pipettes had resistances of 3–5 MΩ. Access resistances were between 20 and 35 MΩ; if access resistance changed >20%, the recording was discarded. Whole-cell voltage-clamp recordings of AMPAR-mediated EPSCs were performed at a holding potential between −60 and −50 mV using an internal solution containing the following (in mM): 60 CsCl, 30 Cs d-gluconate, 20 tetraethylammonium-Cl, 20 EGTA, 4 MgCl2, 4 ATP, and 30 HEPES, pH 7.3, adjusted with CsOH. Paired-pulse experiments were performed at an interpulse interval of 50 ms; the paired-pulse ratio (PPR) was calculated as the ratio between the second and first EPSCs.

### Behavioral Assays

Male *RiboTag::Drd2f/f, RiboTag::Drd2+/+, Drd2-f/f and Drd2-*+/+ mice were injected with AAV8-Cmv-Cre-eGFP and AAVDJ-Pcp2-Cre viruses. Similarly, wild type C57Bl/6j mice were injected with AAV8-Cmv-eGFP and AAV8-Cmv-m-Drd2-IRES-eGFP. Behavioral evaluation was performed 21 days after surgery. Mice were handled for 3 days prior to testing for habituation. All experiments were blinded to genotype during behavioral testing.

#### Locomotor Activity

Horizontal and vertical activity was measured in a circular corridor (Imetronic, Pessac, France) for 60 min. Counts for horizontal activity were incremented by consecutive interruption of two adjacent beams placed at a height of 1 cm per 90° sector of the corridor (mice moving through 1/4 of the circular corridor) and counts for vertical activity (rearings) corresponding to interruption of beams placed at a height of 7.5 cm along the corridor (mice stretching upwards) were used as a measure for exploratory activity. The circular corridor was cleaned with 70% ethanol and dried between each test.

#### Accelerating Rotarod

Accelerating rotarod was used to measure the cerebellar learning as it was previously described^81^. For 2 consecutive days, the mice were trained to hold onto the rod at a constant speed (4 rpm) for at least 60 seconds. On the test day, the rod accelerated from 4 to 40 rpm within 5 minutes and the speed and the latency to fall were recorded over 10 consecutive trials. Data is expressed as: (Latency to fall on trial 10^th^ - Latency to fall on trial 1^st^) and (Speed on trial 10^th^ – Speed on trial 1^st^).

#### Coat-hanger test

We used a coat hanger (diameter: 2 mm, length: 40 cm) divided into 12 segments (length: 5 cm) and suspended at 40 cm from cushioned surface to evaluate the fine motor coordination as it was previously described^81^. Mice were placed in the middle of the hanger, and the fall latency and the number of movements were evaluated for a total of 60 seconds. The test finishes when the animal fells down or when the 60 seconds elapsed.

#### Beam-walking test

Beam-walking test was performed to evaluate the fine motor coordination as it was previously described^82^. Mice were trained to cross a horizontal wooden circle rod (1 m long, 3 cm in diameter) placed 60 cm above a cushioned table for 2 days before the test. The day of the test, mice crossed twice a wide wooden rod (3 cm in diameter) and a narrow wooden rod (1 cm in diameter). The total number of footslips was recorded in each mouse and used to calculate the mean for each experimental group.

#### Three-Chamber Social Approach

A three-chamber arena was used to assess sociability and preference for social novelty similarly as it was previously published^83^. The rectangular arena (60 × 40 × 22 cm) is divided by Plexiglas walls with openings of 8 cm in diameter allowing the access into each chamber (20 × 40 × 22 cm). The middle chamber was used to place the evaluated mouse at the beginning of each trial. The two side chambers contained an empty squared wire cage (8 × 10 cm) in which the social (strangers) or the non-social (object) stimuli could be enclosed. Wire cages present small holes allowing the visual, auditory, olfactory and tactile contact of tested mice with the stimuli confined inside the cages. A weighted cup was placed on the top of the wire cages to prevent the tested mice to climb.

The three-chamber test consisted in three phases of 10 min. In the first phase (Habituation), the evaluated mouse was placed in the middle chamber and allowed to freely explore the three chamber compartments for a period of 10 minutes. During the second phase (Sociability), an unfamiliar juvenile mouse (Stranger1) (from the same mouse line and same gender) that had no prior contact with the tested mouse, was placed in one of the wire cages. In the opposite side chamber, the nonsocial stimuli (Object) was equally enclosed and the evaluated mouse was allowed to freely explore the three chambers for 10 min. During the third face of the test (Preference for Social Novelty), the nonsocial stimuli (Object) was replaced by a new unfamiliar mouse (Stranger 2). The tested mouse was then given the possibility to interact with both stranger 1 and stranger 2 for 10 min. Mice used as social stimuli (Strangers) were habituated to the wire cage for a brief period of time for 3 days before the start of the experiment. Every session was video-tracked and the number of entries that the tested mouse performed in each chamber, the total time it spent in each compartment and in interaction with the object and strangers were manually scored by an experimenter blind to the genotype of the animals. Mice were considered to be exploring the Object and the Strangers when their nose was directed toward the cage at a distance less than approximately 1cm or directly interacting with the wire cages enclosing stimuli. The arena was cleaned with 70% of ethanol and dried during few minutes between trials.

## Statistical analyses

GraphPad Prism v6.0 software was used for statistical analyses. For normally distributed parameters, Student’s t test was used. Multiple comparisons were performed by one-way and two-way ANOVA or two-way repeated measures ANOVA followed by a post-hoc analysis. All data are presented as mean ± SEM, and statistical significance was accepted at 5% level. *p<0.05, **p<0.01, ***p<0.001. For more detailed information about the statistics, check the document “Statistical report by figures”.

## Acknowledgments

We thank Dr. Endo Shogo and Dr. Philippe Isoppe for providing us antibodies against G-substrate and Zebrin-II. This work was supported by Inserm, Fondation pour la Recherche Médicale (DEQ20160334919), La Marató de TV3 Fundació, ANR EPITRACES (EV), ANR DOPAFEAR (EV) and NARSAD Young Investigator Grant from the Brain and Behavior Research Foundation (EP). LC is supported by the post-doctoral Labex EpiGenMed fellowship («Investissements d’avenir» ANR-10-LABX-12-01), EP was a recipient of Marie Curie Intra-European Fellowship IEF327648 and is currently a recipient of Beatriu de Pinós fellowship (# 2017BP00132) from University and Research Grants Management Agency (Government of Catalonia). LC is supported by the pre-doctoral Labex EpiGenMed («Investissements d’avenir» ANR-10-LABX-12-01).

## Author contribution

L.C, E.P and E.V conceived and led the project. L.C, E.P and E.V designed the study. L.C, E.P and E.V conceptualized the project. L.C and E.P performed brain dissections. L.C and E.P performed polysome IP and qRT-PCR experiments. L.C performed western blot analyses. L.C, E.P and P.T performed immunofluorescence assessments. L.C and L.C performed *in situ* hybridization analysis. L.C performed stereotaxic injections and behavioral experiments. L.C, M.B and A.C performed iDISCO methodology. L.C and F.B performed *in vivo* cerebellar slices. L.C and M.A-L designed AAVDJ-Pcp2-Cre, L.C and F.A performed PCs-sorting, M.R provided D2R-floxed mice. M.M performed electrophysiological analyses. E.V supervised the project. L.C and E.V wrote the manuscript with input from all authors. The authors declare no conflicts of interest.

